# A learning-based framework for miRNA-disease association identification using neural networks

**DOI:** 10.1101/276048

**Authors:** Jiajie Peng, Weiwei Hui, Qianqian Li, Bolin Chen, Qinghua Jiang, Xuequn Shang, Zhongyu Wei

## Abstract

**Motivation:** A microRNA (miRNA) is a type of non-coding RNA, which plays important roles in many biological processes. Lots of studies have shown that miRNAs are implicated in human diseases, indicating that miRNAs might be potential biomarkers for various types of diseases. Therefore, it is important to reveal the relationships between miRNAs and diseases/phenotypes.

**Results:** We propose a novel learning-based framework, MDA-CNN, for miRNA-disease association identification. The model first captures richer interaction features between diseases and miRNAs based on a three-layer network with an additional gene layer. Then, it employs an auto-encoder to identify the essential feature combination for each pair of miRNA and disease automatically. Finally, taking the reduced feature representation as input, it uses a convolutional neural network to predict the final label. The evaluation results show that the proposed framework outperforms some state-of-the-art approaches in a large margin on both tasks of miRNA-disease association prediction and miRNA-phenotype association prediction.

**Availability:** The source code and data are available at https://github.com/Issingjessica/MDA-CNN.

**Supplementary information:** Supplementary data are available at *Bioinformatics* online.

## 1 Introduction

A microRNA (miRNA) is a type of non-coding RNA containing about 22 nucleotides and it has been found in different types of organisms, e.g., plants, animals, viruses [Liu *et al.*, 2015, Huang *et al.*, 2011, Li *et al.*, 2013a, Yuan *et al.*, 2015], etc. Existing research reveals that miRNAs play important roles in different biological processes, including cell growth [Ambros, 2003], cell cycle regulation [Carleton *et al.*, 2007], immune reaction [Taganov *et al.*, 2006], tumor invasion [Ma *et al.*, 2007] and cell fate transitions [Shenoy and Blelloch, 2014] by targeting specific messenger RNAs (mRNAs) and regulating gene expression and mRNA degradation [Shi *et al.*, 2013, Jiang *et al.*, 2010].

Studies also indicate that many miRNAs are implicated in human diseases [Jiang *et al.*, 2008], e.g., cancer [Volinia *et al.*, 2012], immune-related diseases [Huang *et al.*, 2007], Parkinson’s disease [Kim *et al.*, 2007], etc. According to the statistics of the Human microRNA Disease Database (HMDD) [Li *et al.*, 2013b], there are more than 10,000 identified miRNA-disease associations, involving 572 miRNAs and 378 diseases. Moreover, recent experiment results [Goulart *et al.*, 2015, Dweep and Gretz, 2015] show strong links between miRNAs and phenotypes. Considering miRNAs might be potential biomarkers for various types of diseases, it is important to further explore the relationships between miRNAs and diseases/phenotypes to understand pathogenicity mechanisms.

Identifying the associations between a pair of miRNA and disease/phenotype via biological experiment is time-consuming and costly in terms of financial input. Therefore, researchers explore to predict these associations automatically based on computational approaches [Pallez *et al.*, 2017, Zou *et al.*, 2015, You *et al.*, 2017]. Most of existing approaches are based on network. The basic assumption is that miRNAs associated with the same or similar diseases are more likely to be functionally related [Chen *et al.*, 2012], and the crucial technical point is similarity computation for different kinds of pairs including miRNA-miRNA, disease-disease and miRNA-disease [Zou *et al.*, 2015]. According to information involved in similarity computation, network-based approaches can be loosely grouped into two categories [Zou *et al.*, 2015], local network similarity methods [Kertesz *et al.*, 2007, Lewis *et al.*, 2003, Wei *et al.*, 2012, Xuan *et al.*, 2013, Han *et al.*, 2014] and global network similarity methods [Chen *et al.*, 2012, Chen and Yan, 2014].

Jiang *et al.* [Jiang *et al.*, 2010] propose a hypergeometric distribution-based method to identify the miRNA-disease association based on miRNA-miRNA and disease-disease network. Xuan *et al.* propose HDMP method to predict miRNA-disease associations based on the most similar neighbors [Xuan *et al.*, 2013]. Specifically, the association score between a miRNA *mr* and a disease *d* is calculated by considering the association between *d* and the *k* most similar neighbors of *mr*. The evaluation results show that HDMP performs better than Jiang’s method. Local network similarity-based methods only consider the direct edge information contained in the involved networks, neglecting the global structure of these networks.

In the global network similarity-based category, some researchers adopt the random walk with restart (RWR) model to predict miRNA-disease associations. For example, RWRMDA utilize RWR on a miRNA-miRNA association network to measure the similarities between known disease-associated miRNAs and candidate miRNAs [Chen *et al.*, 2012]. The experiment results show that RWRMDA obtains reliable predictive accuracy, but it cannot predict novel miRNAs for diseases that do not have known associated miRNAs [Chen and Yan, 2014]. To overcome this drawback, Chen *et al.* propose a method to predict the miRNA-disease association using a semi-supervised method [Chen and Yan, 2014]. A key step of network similarity-based methods is mapping miRNAs and diseases to the same network. Therefore, the result may highly depend on the quality of constructed networks. Recently, You *et al.* construct a heterogeneous network by integrating different types of heterogeneous biological datasets and propose a path-based method to calculate the association score between miRNAs and diseases [You *et al.*, 2017]. Experiment results show that it performs better than other four recently proposed methods ([Xuan *et al.*, 2013], [Chen and Yan, 2014], [Chen *et al.*, 2016] and [Chen *et al.*, 2012]).

Although many attempts have been made to identify miRNA-disease association automatically, most of them are un-supervised ones solely based on networks without using labeled information. This largely limits the performance of state-of-the-art systems. Instead of using networks to compute association scores directly, we explore to use them extract interaction features for miRNA-disease pairs and predict their associations in a supervised fashion.

In general, miRNA-disease networks contain two levels, one for miRNAs and the other for diseases. Similarities are computed within the same level and cross levels. We argue that similarity computation using target concepts (miRNA and disease) directly is unable to catch the deep interaction patterns between them. As we know, miRNAs are implicated in many human diseases by regulating their target genes [Ardekani and Naeini, 2010], indicating that gene plays a crucial role to bridge miRNAs and diseases. For example, miRNA-21 is associated with human hepatocellular cancer via regulating expression of the PTEN tumor suppressor gene [Meng *et al.*, 2007]. Therefore, we believe gene information should be considered in studying miRNA-disease associations. Although some existing methods consider gene information in similarity computation, they can not capture the complex interaction features of how miRNA associating with disease via genes. To address this problem, we introduce a gene layer in the middle of the miRNA and disease layers to form a three-layer-network and propose a novel way to extract interaction features for miRNA-disease pairs.

In past a few years, deep neural networks (e.g. convolution neural networks) have been applied to many bioinformatics applications [Min *et al.*, 2017] and produced promising results. It has been proved that convolutional neural networks obtain the ability to detect meaningful combinations of features from the input data automatically [Angermueller *et al.*, 2016]. Considering features extracted from three-layer network might contain noises, we propose to use convolutional neural network (CNN) to learn the best combination of features and predict the final labels for a given miRNA-disease pair.

In this article, we present a novel learning-based framework, MDA-CNN, to identify the association between a pair of miRNA and disease. Here are four major contributions:

- We introduce a learning-based framework for the task of miRNA-disease association prediction that contains three components, namely network-based feature extractor, auto-encoder-based feature selector and CNN-based association predictor.
- To better represent the correlation between miRNAs and diseases, we construct a three-layer network with an additional gene layer in the middle. A novel feature representation is then proposed based on the regression model.
- We employ a deep convolutional neural network (CNN) architecture to deal with the feature vectors produced in the previous step to determine the final label for miRNA-disease pairs.
- The evaluation results show that MDA-CNN outperforms some state-of-the-art approaches for both miRNA-disease and miRNA-phenotype association identification.

## 2 Methods

We propose a novel algorithm called MDA-CNN to predict miRNA-disease association. The framework of MDA-CNN is shown in figure 1 and it contains three steps. First, given a three-layer network (Figure 1a), we apply a regression model to calculate the disease-gene and miRNA-gene association scores and generate feature vectors for disease and miRNA pairs based on these association scores. Second, given a pair of miRNA and disease, corresponding feature vector is passed through an auto-encoder-based model to obtain a low dimensional representation (Figure 1b). Third, a deep convolutional neural network (CNN) architecture is constructed to predict the association between miRNA and disease based on the representation vector obtained in last step (Figure 1c).

**Fig. 1.**
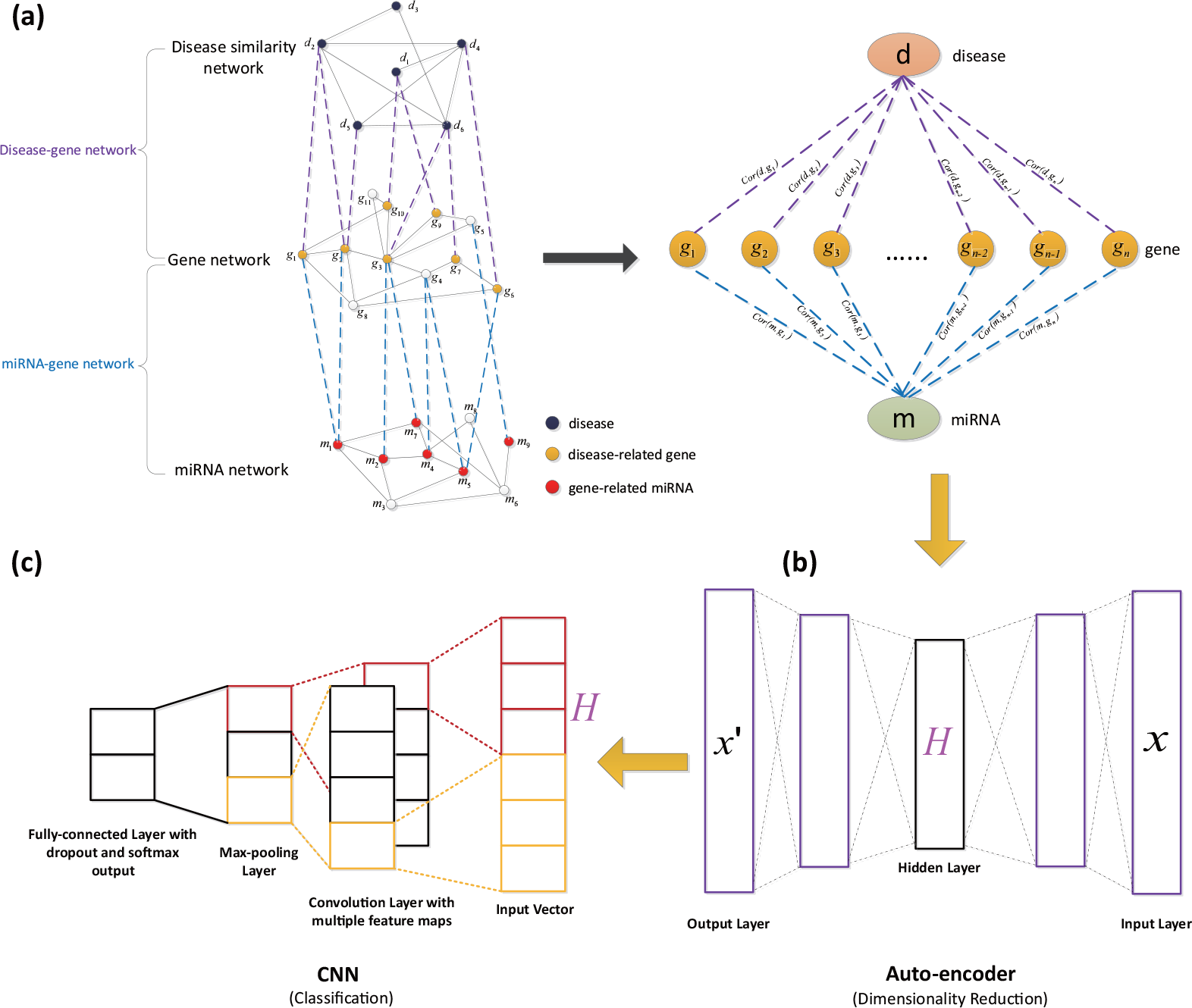
The workflow of MDA-CNN.With a three-layer network as input, MDA-CNN contains three main steps: network-based feature extraction (a), auto-encoder-based feature selection (b) and convolutional neural network-based association prediction (c).

### 2.1 Network-based feature extraction

As we know, miRNAs are implicated in many diseases by regulating gene expression post-transcriptionally. In this work, we add the gene-layer network as the bridge to extract the interaction features between miRNA-disease pairs. For each miRNA-disease pair, its feature vector is the concatenation of the disease vector and miRNA vector. Elements in the disease (or miRNA) vector represent the relations between the disease (or miRNA) and each gene in the gene-layer network. Instead of the binary value that represents whether a disease (or a miRNA) and a gene are associated, we use an association score to measure the relation between a disease (or a miRNA) and a gene.

#### Association score calculation

In the following, we take disease and gene layer as the example to illustrate our algorithm. The association score between a miRNA and a gene can be computed similarly.

Let *N_d_* and *N_g_* be a disease network and a gene association network respectively. 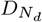 and 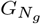 represent the sets of diseases and genes involved in *N_d_* and *N_g_* respectively. *A*_*dg*_ is a set of disease-gene associations between 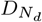 and 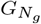. Inspired by [Wu *et al.*, 2008], the association score between a disease *d* and a gene *g* can be measured as the Pearson correlation coefficient of *S_d_* and *R_g_*:

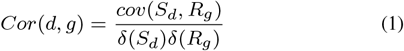

where *S_d_* = [*Sim*(*d*, *d*_1_), *Sim*(*d*_1_, *d*_2_),…, *Sim*(*d*, *d_n_)*] is a vector of similarity scores between *d* and each disease in *N_d_*, *R_g_* = [*R*(*g*, *d*_1_), *R*(*g*, *d*_2_),…, *R*(*g*, *d_n_)*] is a vector of closeness scores between *g* and each disease in *N_d_*, *cov*() and *δ*() represent covariance and standard deviation respectively.

Given *N_g_* and 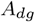, the closeness scores between a gene *g* and a disease *d* can be defined as follows:

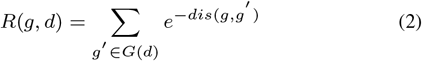

where *G*(*d*) is the set of genes associated with *d*, *dis*(*g*, *g′*) is the square of the shortest path between *g* and *g*′ in *N_g_*.

Instead of using the path-distance-based similarity, we apply a regression model to calculate the similarity between two disease *d_i_* and *d_j_*. The model is able to consider the importance of genes for different diseases. The model is defined as follows:

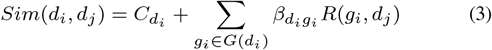

where 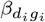 is the regression coefficient of this linear regression model, *G*(*d_i_*) is the set of genes associated with *d_i_*, 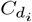 is a bias constant for each disease. 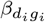 represents the importance of *g_i_* to *d_i_*. The basic idea of this regression model is that the similarity of two diseases can be measured via associated genes. Given *N_d_*, *N_g_* and *A*_*dg*_, this linear regression model could be trained and used to calculate disease similarities. Note that *Sim*(*d_i_*, *d_j_*) can be different with *Sim*(*d_j_*, *d_i_*).

#### Feature representation

Feature representation is the key step before applying the machine learning algorithm. However, existing studies use an association score to link a disease and a miRNA, which is not able to capture the complex interaction between them. By adding the gene layer, we can generate a vector to represent features of a miRNA-disease pair.

Given a disease *d*, we calculate association score between *d* and each gene involved in the gene layer based on Equation 1. After that, a feature vector of *d* can be generated as

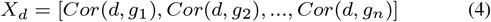

where *g_i_* represents a gene involved in *N_g_*, *n* is the number of genes involved in *N_g_*. To reduce the influence of extreme values (outliers) in *X_d_*, we apply softmax normalization [Grover and Leskovec, 2016] on *X_d_*. Specifically, the normalized vector 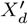 is represented as follows:

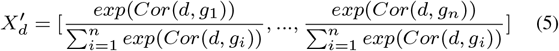

Similarly, given a miRNA *m*, a vector 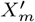 can be generated as follows:

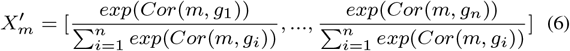

For a miRNA-disease pair, we concatenate 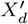 and 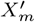 as the vector for feature representation.

### 2.2 Auto-encoder-based feature selection

The length of vector generated in previous step (e.g. the concatenation of 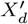 and 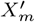) is the double of 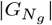, which is very large and noisy. Therefore, we apply auto-encoder to identify the essential feature combination and reduce the dimension of the feature vector for each pair of miRNA and disease automatically.

Auto-encoder is used for dimensionality reduction for the downstream machine learning task such as classification, visualization, communication and the storage of high-dimensional data [Chicco *et al.*, 2014]. Unlike the widely-used method, principal components analysis (PCA), auto-encoder is a non-linear generalization of PCA that uses an adaptive “encoder” network to transform the high-dimensional data into a low-dimensional code, and a similar “decoder” network to recover the data from the lowdimensional code. The low-dimensional code is then used as a compressed representation of the original data.

In our experiments, the vector 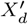 and 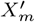 are concatenated before feeding into the auto-encoder model. Let *n* be the number of genes involved in the network *N_g_*. The original dimension of the input is 2*n*. In our model, we use the mean-square error (MSE) [Wax and Ziv, 1977] as the loss function. The sigmoid activation function and Adam algorithm are used to optimize the MSE loss. Our auto-encoder network is trained by the backpropagation (BP) algorithm [Rumelhart *et al.*, 1988].

### 2.3 Convolutional neural network-based association prediction

Convolutional neural network (CNN) was proposed in the late 1980s by Lecun [Lecun *et al.*, 1989] which performs very well in image classification [Krizhevsky *et al.*, 2012], sentence classification [Kim, 2014] and classification task on the structured-graph data [Atwood and Towsley, 2015]. In this work, we also choose convolutional neural network as supervised learning model for learning the best combination of features and predicting the final label of a given miRNA-disease pair. The structure of the proposed model is shown in Figure 2. Our model includes the following layers: convolution and activation layer, max-pooling layer, fully-connected layer and softmax layer. A convolutional layer and a rectified linear unit (ReLU, [Nair and Hinton, 2010]) activation layer are used to extract features from the input that is the output of dimension reduction step (see previous subsection). A pooling layer is used for dimensionality reduction. The final fully-connected layer and softmax layer are for the classification task.

**Fig. 2.**
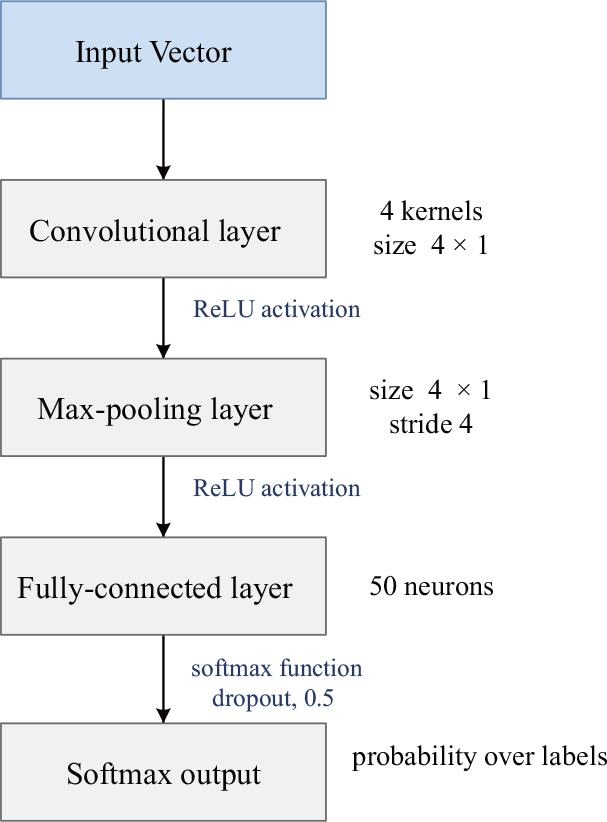
The structure of the proposed convolutional neural network model. The input is a vector. Our model consists of the convolutional layer, max-pooling layer and FC layers. The output is the probability distribution over labels for each sample.

The convolutional layer is responsible for learning subspace features of the input. The convolutional layer of our model consists of 4 kernels. 4 × 1 weight vector is convolved with the input vector with length *L*. After the convolution, for each kernel, we can obtain a feature map *C* (a particular feature extracted from the input) that is a vector with the length (*L* − 4) + 1. The feature map *C* is extracted by the following equation:

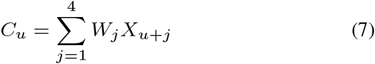

where *u* ∈ {0,…, *L* − 4}, *X* is the input vector, and *W* ∈ *R*^4^ is a weight vector, which is initialized by a truncated normal distribution with mean zero and standard deviation 0.1. A high *C_u_* indicates that the kernel captures the features of the sub-region of the input very well. *C_u_* is then passed through a ReLU function *f*(*x*) = *max*(0, *x*) that ignores negative outputs and propagates positive outputs from previous layer. Although there are various non-linearities, ReLU activations are the most popular because of its computational efficiency, sparsity and reduced likelihood of vanishing gradient [Krizhevsky *et al.*, 2012, Lecun *et al.*, 2015].

Max-pooling layer is used to down-sample the latent representation after the convolutional layer. It takes the maximum value over nonoverlapping sub-regions (i.e., the pooling size is 4) of the output of convolutional layer and outputs the most important feature over a neighborhood in each feature map. Given an input sequence *f*(*C_u_*) (*u* ∈ {0,…, *L* − 4}), the output of pooling layer is shown as follows:

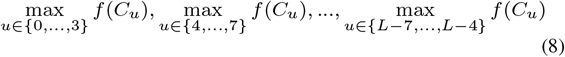

with length of 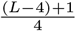.

Convolutional layer and max-pooling layer can extract the important features from the input vector. The outputs of all the kernels are then concatenated into a vector and fed to the fully-connected layer.

The final two layers are a fully-connected layer and a softmax layer. There are 50 hidden units in the fully-connected layer. The output of pooling layer is *y* ∈ *R^n^*, where *n* is the length of concatenated outputs of the pooling layer. The output of the fully-connected layer is :*f* (*W* · *y*), where *W* ∈ *R*^50×*n*^ is the weight matrix, and *f* is the ReLU activation. The final softmax layer is used for classification task.

## 3 Results

### 3.1 Experiment setup

We evaluate our model on two tasks, namely, miRNA-disease association prediction and miRNA-phenotype association prediction.

For miRNA-disease association prediction, we need to have three similarity networks for the same type of elements, namely, diseases, genes and miRNAs. We obtain disease similarity network and miRNA similarity network from You *et al.* [2017]. We utilize the protein network of human genes from Human Protein Reference Database (HPRD) [Baolin and Bo, 2007]. The involved associations across different networks are disease-gene and miRNA-gene associations. The disease-gene associations are obtained from DisGeNET database [Piñero *et al.*, 2016] and only manually curated disease-gene associations are kept. The miRNA-gene associations are obtained from miRWalk2.0 database [Dweep and Gretz, 2015]. During the computation, we remove those genes that have no correlation with diseases or miRNAs. For the experimental dataset, the positive set is obtained from HMDD [Li *et al.*, 2013b]. Because there is no available dataset for negative samples, we randomly generate a negative set with the same size as the positive set.

In the evaluation of miRNA-phenotype association prediction, the three involved networks are phenotype, protein and miRNA network. The protein and miRNA networks are the same as the dataset used in the miRNA-disease association prediction task. The phenotype network is generated based on HPO [Robinson *et al.*, 2008]-based similarity using traditional Resnik method [Resnik, 1995], which is also utilized in [Masino *et al.*, 2014]. The involved associations across networks at different levels are phenotype-gene and miRNA-gene associations. The miRNA-gene associations are the same as the dataset used in the miRNA-disease association prediction test. Phenotype-gene associations are obtained from HPO database. For performance evaluation, the positive set of validated miRNA-phenotype associations is obtained from miRWalk2.0 database [Dweep and Gretz, 2015]. We randomly generate a negative set with the same size as the positive set.

In both tasks, we use 10-fold cross validation [Kohavi *et al.*, 1995] to train and test our model. All aforementioned datasets are available in the supplementary data. The detail of these datasets could be found in supplementary document. The parameters used in the auto-encoder and convolutional neural network are also described in the supplementary document.

### 3.2 Performance evaluation on predicting miRNA-disease associations

We evaluate the performance of MDA-CNN and the other three methods (i.e. CIPHER [Wu *et al.*, 2008],PBMDA [You *et al.*, 2017] and WPSMDA) on the task of predicting miRNA-disease associations. WPSMDA is an alternative of MDA-CNN. The detail of WPSMDA can be found in the supplementary document. Evaluation metrics include area under the receiver operating characteristic curve (AUROC), area under the precision-recall curve (AUPR), precision, recall and F1-score.

The experiment results show that MDA-CNN achieves the highest performance among all methods according to AUROC, AUPR, and F1-Score. The average AUROC achieved by MDA-CNN across 10-fold cross validation is 0.8897, which is significantly higher than the scores of CIPHER, PBMDA and WPSMDA (Table 1). Figure 3(a) shows ROC curves of one cross-validation result. Results of other 9 splits can be found in the supplementary document (Supplementary Figure S2). Comparing the AUPR scores from the four methods shows that MDA-CNN performs the best with the PBMDA as the runner-up (Table 1 and Figure 3(b)). The average AUPR score of MDA-CNN is about 0.27 higher than the second best method. Figure 3(b) shows P-R curves of one cross-validation result. Results of other 9 splits can be found in the supplementary document (Supplementary Figure S3). It is shown that MDA-CNN can achieve the highest F1-score. Importantly, both precision and recall scores of MDA-CNN are more than 0.80 when achieving the highest F1-score. In summary, this experiment shows that MDA-CNN can achieve significant improvement in predicting miRNA-disease associations compared to some state-of-the-art approaches.

**Table 1.** The AUROC, AUPR, Precision, Recall and F1-Scores of four methods on miRNA-disease association prediction task. Bolded numbers are the best performance in each category.

|  | AUROC | AUPR | Precision | Recall | F1-score |
| --- | --- | --- | --- | --- | --- |
| CIPHER | 0.5564 | 0.5612 | 0.4942 | **0.9954** | 0.6605 |
| PBMDA | 0.6321 | 0.6140 | 0.5192 | 0.9036 | 0.6594 |
| WPSMDA | 0.6406 | 0.5942 | 0.5418 | 0.9363 | 0.6864 |
| MDA-CNN | **0.8897** | **0.8887** | **0.8244** | 0.8056 | **0.8144** |

**Fig. 3.**
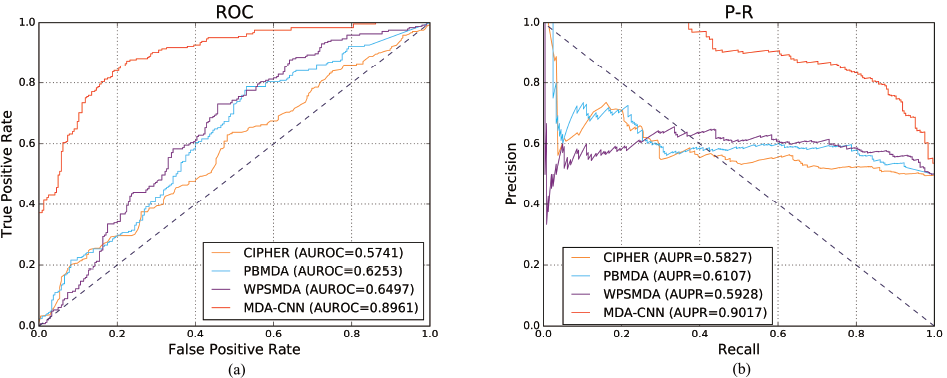
The ROC (a) and P-R (b) curves of four tested methods on miRNA-disease association prediction task.

### 3.3 Performance evaluation on predicting miRNA-phenotype associations

In addition to evaluate MDA-CNN on miRNA-disease association prediction problem, we further test whether MDA-CNN can be applied to predict miRNA-phenotype associations. Patient phenotypes, which may be determined by both genetically or environmentally, are the physical, biochemical and physiological makeup of a patient [Robinson *et al.*, 2008]. Understanding the miRNA-phenotype association can reveal how miRNAs are implicated in human diseases.

Similar with the evaluation on miRNA-disease dataset, we compare MDA-CNN with three methods (i.e. CIPHER, PBMDA and WPSMDA).In general, the results show that MDA-CNN performs better than other methods according to all metrics.The average AUROC score of MDA-CNN is 0.9429, which is significantly higher than the second best method PBMDA (the value is 0.7453) (Table 2). Similarly, MDA-CNN achieves the highest AUPR score (0.9344), while the score of the runner-up (PBMDA) is 0.7197 (Table 2). Figure 4(a) and (b) show ROC and PR curves of one cross-validation result. Results of other 9 splits can be found in supplementary document (Supplementary Figure S4 and S5). It is shown that MDA-CNN can achieve the highest F1-score (Table 2). Comparing with other methods, MDA-CNN can achieve both high precision and recall. In summary, MDA-CNN performs better than some state-of-the-art approaches in predicting miRNA-phenotype associations.

**Table 2.** The AUROC, AUPR, Precision, Recall and F1-Scores of four methods on miRNA-phenotype association prediction task. Bolded numbers are the best performance in each category.

|  | AUROC | AUPR | Precision | Recall | F1-score |
| --- | --- | --- | --- | --- | --- |
| CIPHER | 0.5212 | 0.5271 | 0.5051 | **0.9991** | 0.6710 |
| PBMDA | 0.7453 | 0.7197 | 0.6725 | 0.7614 | 0.7141 |
| WPSMDA | 0.6927 | 0.6954 | 0.5492 | 0.9310 | 0.6909 |
| MDA-CNN | **0.9429** | **0.9344** | **0.8667** | 0.8748 | **0.8704** |

**Fig. 4.**
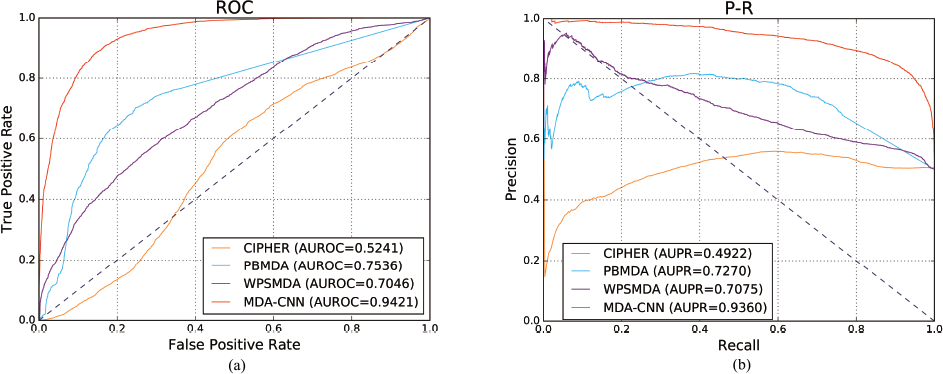
The ROC (a) and P-R (b) curves of four tested methods on miRNA-phenotype association prediction task.

### 3.4 Effects of MDA-CNN components

In order to evaluate the performance of each step of MDA-CNN, we compare MDA-CNN with two versions of MDA-CNN, each with a different approach in feature representation step and convolutional neural network step on miRNA-disease association prediction task. To test the performance of our feature representation model, we create BR-CNN where the values in the feature vector are binary values. To test the effect of the convolutional neural network model, we create MDA-SVM where the convolutional neural network model is replaced by support vector machine (SVM) model. Table 3 shows that MDA-CNN is significantly better than BR-CNN (p-value < 0.05, Wilcoxon test) and MDA-SVM (p-value < 0.05, Wilcoxon test) in terms of all metrics, indicating that the two steps in MDA-CNN contribute to the performance and have been appropriately designed.

**Table 3.** The AUROC, AUPR, Precision, Recall and F1-Scores of MDA-SVM, BR-CNN and MDA-CNN on miRNA-disease association prediction task. Bolded numbers are the best performance in each category.

|  | AUROC | AUPR | Precision | Recall | F1-score |
| --- | --- | --- | --- | --- | --- |
| MDA-SVM | 0.8647 | 0.8524 | 0.7879 | 0.7856 | 0.7864 |
| BR-CNN | 0.8537 | 0.8515 | 0.7884 | 0.7726 | 0.7797 |
| MDA-CNN | **0.8897** | **0.8887** | **0.8244** | **0.8056** | **0.8144** |

### 3.5 Case study on lung cancer

We apply MDA-CNN to predict the miRNAs associated with lung cancer as a case study.

Lung cancer is the leading cause of cancer-associated death [Bandi *et al.*, 2009, Jemal *et al.*, 2010]. In HMDD database, there are 110 miRNAs that associate with lung cancer. In the experiment, to avoid logic circle, we first remove all the associations between lung cancer and its related miRNAs from the training set. We use MDA-CNN to predict the association between lung cancer and every miRNA. Then, we compare the prediction result with the record in HMDD database. 110 miRNAs associating with lung cancer are found in HMDD database. 52 of 110 miRNAs, named “easy set”, have at least one known target gene in the protein network. 58 of 110 miRNAs, named “hard set”, have no known target gene in the protein network. It is no surprise that almost all miRNAs (51 of 52) in the “easy set” are identified by MDA-CNN. In addition, 53 of 58 miRNAs in “hard set” are identified. To test the power of gene layer-based feature representation, we replace gene layer-based feature representation with binary values representing whether a miRNA targets a gene. In the “easy set”, 41 of 52 miRNAs are identified. However, in the “hard set”, only 12 of 58 miRNAs are identified. It indicates that by introducing a gene layer network to extract interaction features, MDA-CNN is able to enhance the performance of miRNA-disease association prediction.

To test whether MDA-CNN can predict new miRNA-disease associations that is not available in existing database. We rank the predicted miRNAs based on their prediction probabilities. We find that 3 of top 20 miRNAs are not associated with lung cancer (see supplementary table S1). The three miRNAs are hsa-mir-16, hsa-mir-15a and hsa-mir-106b, which are in the third, fourth and tenth place in the predicted miRNAs. Based on the literature study, we find that hsa-mir-15a and hsa-mir-16 are likely contributing to the tumorigenesis of non-small cell lung cancer by regulating cyclins D1, D2, and E1 [Bandi *et al.*, 2009]. Although no direct evidence shows that hsa-mir-106b is associated with lung cancer, evidence shows that hsa-mir-106b is a cancer-associated miRNA [Nagini, 2012].

## 4 Conclusion

Recently, researchers have started to focus on identifying miRNA-disease associations by computational tools. In this article, we propose a learning-based framework named MDA-CNN to identify the miRNA-disease/phenotype associations. We first extract features of miRNA and disease/phenotype based on a three-layer network. Then, an autoencoder-based model is proposed for feature selection. Using this feature representation, we propose a convolutional neural network architecture for the purpose of predicting miRNA-disease/phenotype associations. To demonstrate the advantages of MDA-CNN, we compare it with three state-of-the-art methods. The experiments on both miRNA-disease and miRNA-phenotype associations show that MDA-CNN performs better than existing methods, indicating that the proposed learning-based framework is appropriately designed. Additionally, case study on lung cancer shows that MDA-CNN could be used to predict the miRNA-disease associations.

